# *Cryptococcus neoformans* responds to presence of mycobacterium by diversifying its morphologies and remodelling its capsular material

**DOI:** 10.1101/2025.11.16.687457

**Authors:** Orlando Ross, Liliane Mukaremera, Ivy M. Dambuza

## Abstract

*Cryptococcus neoformans* and *Mycobacterium tuberculosis* are opportunistic pathogens that share overlapping geographical distributions and physiological niches within the human body. Both are recognised by the World Health Organization as high-priority pathogens. Although clinical reports of co-infections with cryptococcosis and tuberculosis are increasing, experimental studies exploring their interactions remain scarce. Here, we demonstrate that *C. neoformans* can grow in physiologically relevant human plasma-like medium in the presence of either heat-killed *M. tuberculosis* antigen or the live vaccine strain, *M. bovis* BCG. In response to presence of mycobacteria, *C. neoformans* increased in number and exhibited enhanced virulence-associated traits, including titan cell formation, capsule enlargement and increased survival from phagocytosis. This work provides proof-of-principle for a dynamic, inter-pathogen interaction that may contribute to exacerbation disease outcomes in settings of a co-infection.

## Introduction

*C. neoformans* colonises the pulmonary region through inhalation of spores and desiccated yeast cells, often found in bird guano and hollows of various tree species, including *Eucalyptus* spp., in the environment^1–3^. Pulmonary cryptococcosis results in the formation of nodules or masses in the lung, known as granulomas. In parallel, the pathogenesis of tuberculosis has a closely related cycle; aerosolised *Mycobacterium tuberculosis* (Mtb) droplets are inhaled, reaching the lungs whereby colonisation and granuloma formation also occurs^4,5^. Cryptococcal granulomas are often indistinguishable from those formed by Mtb and impairment of the immune system, such as in HIV/AIDS, can lead to failure to restrain growth of these pathogens within the lung resulting in spread to other organs, including the central nervous system (CNS). Both *C. neoformans* and Mtb CNS infection can result in life-threatening meningoencephalitis if not promptly treated^6,7^. While mechanisms of dissemination to extrapulmonary regions remains elusive, in both cryptococcosis and tuberculosis, it is thought that the pathogens seed distal tissues through trafficking by infected macrophages or through systemic infection when granulomas fail to contain pathogen growth^6,8–13^.

The World Health Organisation (WHO) classified *C. neoformans* as a ‘critical priority’ in the first ever ‘Fungal Pathogens Priority List’ in October 2022^14^. *C. neoformans* is estimated to be implicated in 194,000 cases of HIV-associated cryptococcal meningitis globally, with 118,000 *C. neoformans*-mediated meningitis (CM) related deaths^15,16^. A disease-linked mortality is estimated to be as high as 70% in low-income countries, compared to 20-30% in high-income countries^15,16^. Early-stage diagnosis of cryptococcosis is crucial for reducing mortality, but detection through rapid testing, or affordable methods that do not use PCR/RNA sequencing, are usually only employed when disease is presenting, a sign of advanced CM and often too late^17^. Similarly, the mortality rates caused by Mtb are equally devasting. For instance, Mtb was the leading infectious disease single-agent cause of death globally, before the COVID-19 pandemic, with 50% mortality observed in untreated patients^18,19^. Mtb is estimated to latently infect nearly one quarter of the global population, up to 1.7 billion people^18,20^. The 2022 WHO global tuberculosis report estimated that 10.6 million people became ill and 1.6 million people died from tuberculosis in 2021 alone, 11.7% of deaths occurring in HIV-positive patients^18^. Multi-drug-resistant tuberculosis (MDR-TB) incidence has also risen, further driving morbidity and mortality, alongside healthcare-associated costs^21^.

Yet, mounting evidence demonstrates the increasing occurrence of *Mycobacterium* and *Cryptococcus* spp., co-infections, which is of great public-health concern, especially in sub-Saharan Africa and Asia in both HIV-positive and HIV-negative populations^22–26^. *Mycobacterium* and *Cryptococcus* spp., co-infections significantly increase the risk of death compared to *Cryptococcus* deaths alone^22^. Despite this, we have very little understanding whether these pathogens interact with one another or whether pathogenesis or virulence is increased or hindered in a co-infection setting. We hypothesised that *C. neoformans* would be influenced by the presence of *Mycobacterium* spp., potentially upregulating virulence factors such as cell body size and capsule, as we have previously shown that bacterial cell wall components drive *C. neoformans* cell enlargement^27^. Other fungal-bacterial co-infection settings result in drastic prognostic changes, and confer worse clinical outcomes, such as *Candida albicans-Staphylococcus aureus* co-infections which have devastating consequences for the host^28^. In this study, we demonstrate that *C. neoformans* senses mycobacteria and shifts toward a more virulent state, increasing proliferation and inducing capsule and titan formation. This establishes proof-of-principle that inter-pathogen interactions may potentiate *C. neoformans* pathogenicity during co-infection settings.

## Results

### *C. neoformans* cell body, capsule size and cell density are significantly increased by co-culture with *Mycobacterium* species

We have shown recently that *C. neoformans* produces *in vivo* morphologies when grown *in vitro* using human-plasma like medium (HLPM), 5% CO_2_ and 37 °C^29^. These culture conditions provided ideal *in vitro* settings to test whether *C. neoformans* morphological switching, which is a characteristic virulence factor^30–32^, was impacted by presence of mycobacteria. To assess this, we co-cultured H99 with live *M. bovis* BGC (**Figure 1A**). When we assessed the overall mean cell body size within the cultures, in the presence of BCG at either 24 hours (**Figure 1B and D**) or 48 hours (**Figure 1C and E**) we found no significant change compared with H99 monoculture. However, analysis of size distributions at 24 hours revealed a shift in population structure: the proportions of titan cells (≥10 μm) and yeast-sized cells (5 to 9 μm) increased by approximately 4.5% and 10%, respectively, while the fraction of smaller cells (≤5 μm) decreased by ∼14% (**Figure 1D**). This pattern suggests that H99 senses the presence of live BCG and responds by diversifying its population heterogeneity, generating a higher proportion of yeast-sized and titan morphotypes while reducing small-cell abundance. This response was still detectable after 48 hours of co-culture, although more modestly, with titan and yeast-sized cells increasing by ∼1% and 8%, respectively, and the proportion of smaller cells decreasing by ∼9% (**Figure 1E**). These findings indicate that the interaction between *C. neoformans* H99 and live BCG promotes sustained remodelling of the fungal population structure, consistent with an adaptive sensing mechanism rather than passive morphological drift. We also examined relative to the avirulent BCG, how H99 responds to a virulent mycobacterium strain, Mtb. The absence of a Biosafety level 3 (BCL-3) facilities prevented the use of live Mtb; therefore, heat-killed Mtb (HK-Mtb) was used, which retains complex immunogenic cell-wall lipids and glycolipids associated with its virulence. Notably, co-culture of H99 with HK-Mtb resulted in a statistically significant increase in the mean cell body size of the overall H99 population at both 24 hours (**Figure 1B and D**) and 48 hours (**Figure 1C and E**). Analysis of population size distributions revealed a striking shift at 24 hours, with the proportion of titan cells (≥10 μm) increasing by 27.4%, accompanied by a 13% reduction in yeast-sized cells (5 to 9 μm) and a 14% decrease in smaller cells (≤5 μm) (**Figure 1D**). This pattern was still detectable at 48 hours, although to a lesser extent, with titan cells increasing by 7% and yeast-sized and smaller cells decreasing by 1% and 6%, respectively (**Figure 1E**). These findings indicate that exposure to virulent Mtb or the increased Mtb-associated molecules exposed due to heat killing, resulted in pronounced and sustained restructuring of H99 population morphology, strongly favouring titan cell formation. Importantly, when we assessed the total cell counts, we observed a significant increase in H99 cell density during HK-Mtb co-culture (**Figure 1F**), suggesting that this morphological shift occurs in parallel with overall population expansion rather than growth suppression.

**Figure 1.**
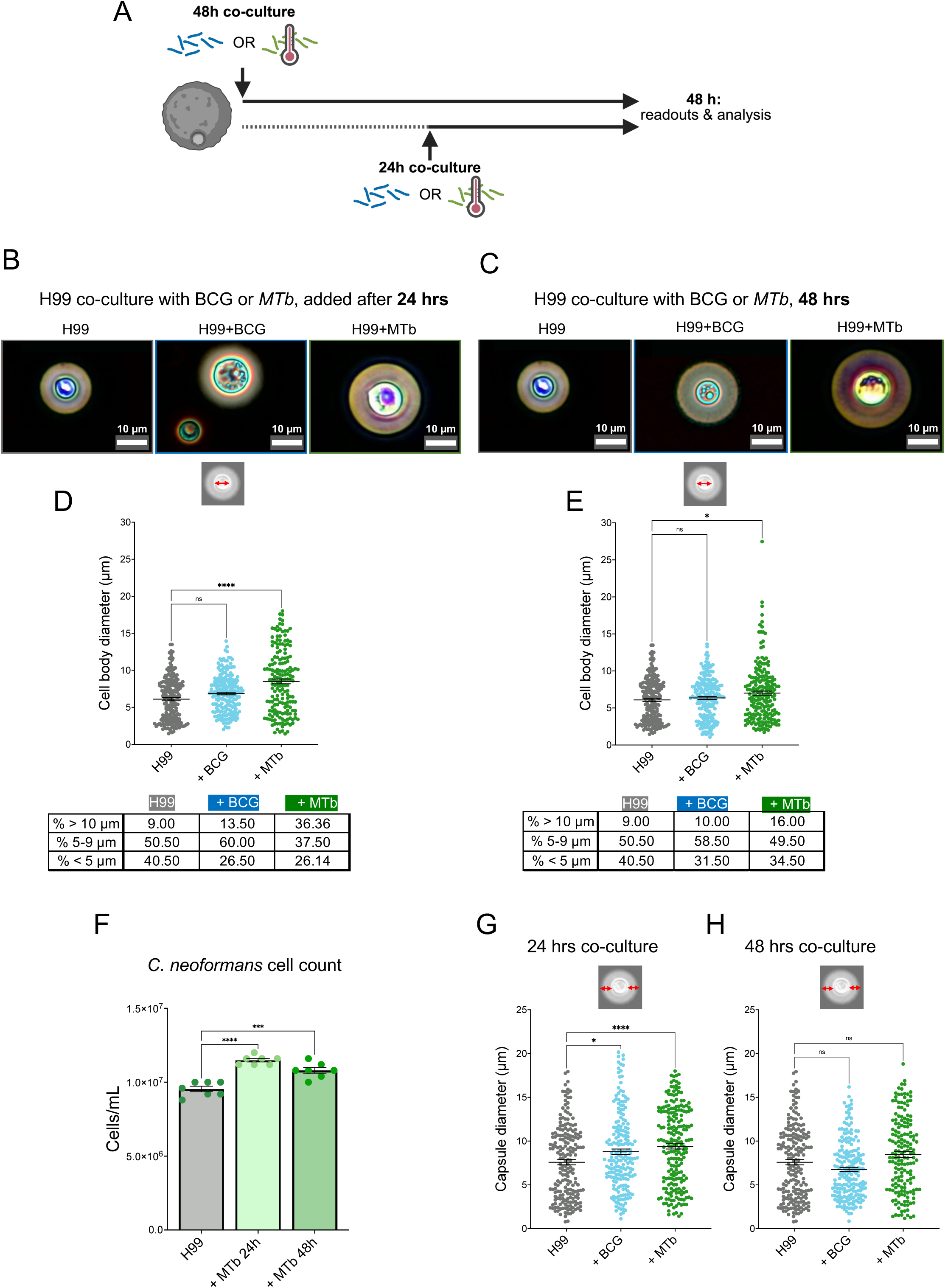
*Cryptococcus neoformans* morphologies are significantly altered in the presence of *Mycobacterium* spp. (**A**) Schematic of culture conditions. *C. neoformans* H99 overnight cells incubated in HPLM supplemented with 10% FBS at 37°C, 5% CO2; either alone (1×10^4^ cells/well) or with *M. bovis* (1×10^3^ cells) or *M. tuberculosis* (2.5 ng heat-killed desiccated H37Ra) for 24h (**B**) or 48h co-culture (**C**). H99 cells were counterstained with india ink, imaged on Olympus EP50 camera, mounted to Olympus CKX53 microscope. **B** and **C** show representative images of cell body diameter and percentages of different subpopulations (> 10 µm, between 5-9 µm and < 5 µm) at 24h (**D**) and (**E**). (**F**) Shows cell density after co-incubation with *M. tuberculosis* quantified using Vi-cell Blu viability analyser. **G** (24h) and **H** (48h) show capsule size (diameter). Data presented are mean ± SEM from 8 technical repeats across 2 biological replicates. All cell measurements were made using Fiji, graphed on GraphPad Prism, n = 200 cells (8 technical repeats across two biological repeats). Tukey’s multiple comparisons test was used to assess statistical significance of populations as a whole; * denotes p ≤ 0.05, *** denotes p ≤ 0.001, **** denotes p ≤ 0.0001, ‘ns’ denotes not statistically significant. FBS: foetal bovine serum, HPLM: human plasma-like media, YPD: yeast peptone dextrose.

The polysaccharide capsule is a well-studied virulence determinant of *C. neoformans*, functioning both as a physical barrier and as a dynamic regulator of host-pathogen interactions^30^. Titan cells are characterized by dramatic capsule thickening associated with enhanced immune evasion and persistence in tissues^31,33^. In our previous work, we established that HLPM, 5% CO_2_ and 37° C *in vitro* conditions reliably induced large capsule formation, closely mirroring capsule expansion observed *in vivo*, and importantly, this phenotype extended beyond titan cells to include smaller cell populations^29^. Given that both cell body enlargement and capsule remodelling contribute to pathogenic fitness, we next examined whether exposure to live BCG or HK-Mtb alters capsule size in H99 populations. Capsule measurements at 24 hours revealed no significant difference in capsule diameter when H99 was co-cultured with either live BCG or HK-Mtb compared with H99 monoculture (**Figure 1G**). However, by 48 hours, both live BCG and HK-Mtb induced a significant increase in capsule thickness (**Figure 1H**). This indicates that prolonged exposure to mycobacterial cues promotes capsule remodelling. Together, these data show that *C. neoformans* H99 expands its capsule in response to either attenuated or virulent mycobacterial components, suggesting that capsule remodelling is a generalized response to mycobacterial sensing rather than a virulence-specific phenomenon.

### Clinical isolates of *C. neoformans* remodel the cell body population and capsule production in presence of live *M. bovis*

Recent work shows that clinical isolates of *C. neoformans* modulate different disease outcomes in patients^34–36^. H99 originates from a Hodgkin’s lymphoma patient, existing in the VNI *C. neoformans* clade^37^ and is not closely related to the majority of strains that are associated with HIV – the key group of patients at risk from *C. neoformans* – Mtb co-infections^34,38^. To determine whether the morphological response to mycobacteria was conserved across diverse *C. neoformans* backgrounds, we examined two high-virulence clinical isolates (SACl012 and UgCl387) and two low-virulence isolates (UgCl223 and UgCl425)^34^ (**Supplementary Figure 1**). When co-cultured with live BCG under HLPM conditions, both SACl012 (**Figure 2A and B**) and UgCl387 (**Figure 2A and C**) exhibited clear increases in mean cell body size and restructuring of population architecture compared with monocultures. SACl012, which does not generate titan cells under these culture conditions, responded by markedly enriching its yeast-sized (5 and 9 μm) population, showing a ∼35% increase in yeasts and a 35% reduction in smaller cells (≤5 μm) (**Figure 2B**). UgCl387, by contrast, not only expanded its yeast-sized population by ∼12% but also generated titan cells *de novo*, with a ∼7% increase in cells ≥10 μm and a concurrent ∼19% decrease in smaller forms (**Figure 2C**). These findings parallel the remodelling observed in H99 and indicate that high-virulence isolates readily diversify their morphological repertoire in response to mycobacterial cues, shifting toward cell types linked to persistence and immune resistance.

**Figure 2.**
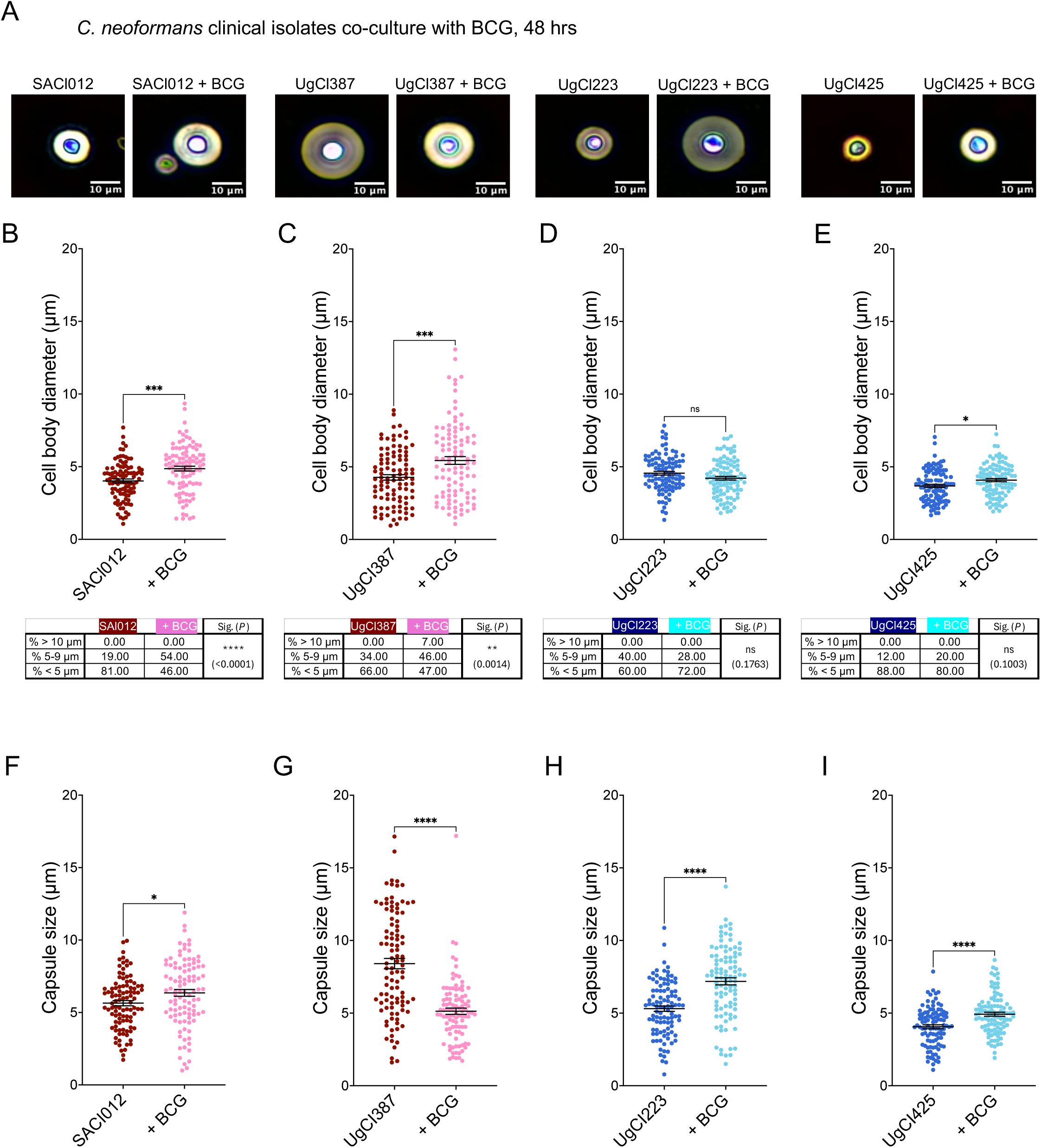
Clinical isolates of *C. neoformans* remodel the cell body population and capsule production in presence of live *M. bovis*. High (in red) and low (in blue) virulence *C. neoformans* clinical isolates were grown overnight in YPD at 30°C with shaking, washed with PBS and then resuspended in HPLM supplemented with FBS in 12-well plates (1×10^4^ cells per well). These cultures were then incubated with or without *Mycobacterium bovis* (BCG) (1×10^3^ cells/well) in HPLM supplemented with 10% FBS at 37°C, 5% CO2 for 48 hours at 37°C and 5% CO2. Cells were counterstained with India ink and observed on Olympus CKX53 microscope, with Olympus EP50 camera. (**A**) shows representative images of India ink-stained cells. (**B** to **E)** show overall percentage cell body size, and the table below each graph indicates the percentages of different *C. neoformans* subpopulation (> 10 µm, between 5-9 µm and < 5 µm). (**F** to **I)** show capsule diameter measured using Fiji. Data presented are from 2 biological replicates with at least 100 cells per replicates. Median with interquartile range shown by error bars. Mann-Whitney U test was used to measure statistical significance between *C. neoformans* strains alone and in co-culture with BCG, * denotes p ≤ 0.05, ** denotes p ≤ 0.01, **** denotes p ≤ 0.0001.

The low-virulence isolates exhibited distinct responses that were not defined simply by attenuated versions of the high-virulence pattern. UgCl223 showed no significant change in mean cell size overall (**Figure 2D**). However, analysis of size distributions revealed a notable increase in smaller-cell forms (≤5 μm) accompanied by a decrease in yeast-sized cells, by ∼12% (**Figure 2D**). This redistribution is not indicative of a neutral or diminished response but instead suggests a shift toward a morphology associated with improved dissemination^39^. Small-cell morphotypes have been reported to traverse endothelial barriers more efficiently and seed distant tissues^39^, implying that, in this isolate, mycobacterial sensing may promote a spread-oriented pathogenic strategy rather than immune evasion through size enlargement. UgCl425, in contrast, displayed a statistically significant increase in mean cell size (**Figure 2E**), driven by an ∼8% expansion of yeast-sized cells and a reduction in smaller forms, without titan cell emergence, representing a more moderate remodelling profile. Taken together, these findings demonstrate that *C. neoformans* does not adopt a uniform response to mycobacterial co-culture. Instead, each isolate shifts along a morphological trajectory that aligns with its underlying virulence program. High-virulence isolates preferentially enriched titan and yeast morphotypes associated with immune evasion and tissue persistence, whereas low-virulence isolates either enhance small-cell production to favour dissemination or expand yeast-sized populations without generating titan forms. Thus, exposure to mycobacteria does not impose a single pathogenic state but rather acts as an environmental signal that amplifies strain-specific virulence strategies already embedded within the genetic identity of each isolate.

We next assessed whether clinical *C. neoformans* isolates remodel their capsule when co-cultured with mycobacteria. All isolates were grown in HPLM and co-cultured with live BCG for 48 hours under host-like conditions. The high-virulence isolate, SACl012, exhibited a significant increase in capsule thickness in response to BCG (**Figure 2F**), mirroring the capsule expansion observed in H99. In contrast, the second high-virulence isolate UgCl387 demonstrated a significantly reduced capsule when exposed to BCG (**Figure 2G**), indicating that even among isolates with similar clinical severity profiles, capsule remodelling can occur in opposing directions. For the low-virulence isolates, both UgCl223 and UgCl425 showed significant capsule enlargement following BCG co-culture compared to the monocultures (**Figure 2H-I**). Notably, these responses occurred despite distinct effects of BCG on cell body size and population structure, suggesting that capsule remodelling is a robust and conserved response to mycobacterial cues, whereas the direction and morphological context of that remodelling differ between isolates. Together, these findings demonstrate that exposure to mycobacteria triggers capsule restructuring across genetically and clinically diverse *C. neoformans* isolates. Rather than a uniform program, *C. neoformans* appears to deploy isolate-specific remodelling strategies, with some isolates increasing capsule thickness and others reducing it. Because capsule architecture is a key determinant of immune evasion, dissemination, and persistence, such plasticity has important implications for co-infection.

### Pre-activation of alveolar-like macrophages with IFN-γ and *M. tuberculosis* cues reduces their capacity to restrict *C. neoformans* growth

Pulmonary alveolar macrophages are among the first immune cells to encounter *C. neoformans* in the lung. To determine whether the morphological and population-level changes induced by mycobacterial exposure alter macrophage-fungal interactions, we next assessed the ability of alveolar-like macrophages (AMs) to restrict *C. neoformans* growth. AMs were generated from murine bone marrow cells by cytokine differentiation^40^ (**Supplementary Figure 2A and B**), and flow cytometric analysis confirmed acquisition of a CD11b^high^, CD11c^+^, Siglec-F^+^, F4/80^high^ phenotype characteristic of resident AMs (**Supplementary Figure 2C**). AMs were pre-stimulated with heat-killed HK-Mtb and recombinant IFN-γ^41–43^, to mimic a type-1 immune environment shaped by presence of Mtb infection, which are known to confer anti-microbial activity to macrophages^44,45^. AM morphology was monitored throughout co-culture (**Figure. 3A-C**, **Supplementary Fig. 2D**). After 18 hours, significantly higher numbers of viable *C. neoformans* cells were recovered from AMs pre-stimulated with IFN-γ and HK-Mtb compared with unstimulated AMs (**Fig. 3D-E**). These data show that AMs conditioned with mycobacterial signals are less effective at restricting *C. neoformans* growth. In the context of the morphological remodelling and increased capsule expansion in response to mycobacterial sensing, this result indicates that *C. neoformans* exposed to mycobacterial environments can persist and expand even within macrophages that display a type-1, classically activated phenotype.

**Figure 3.**
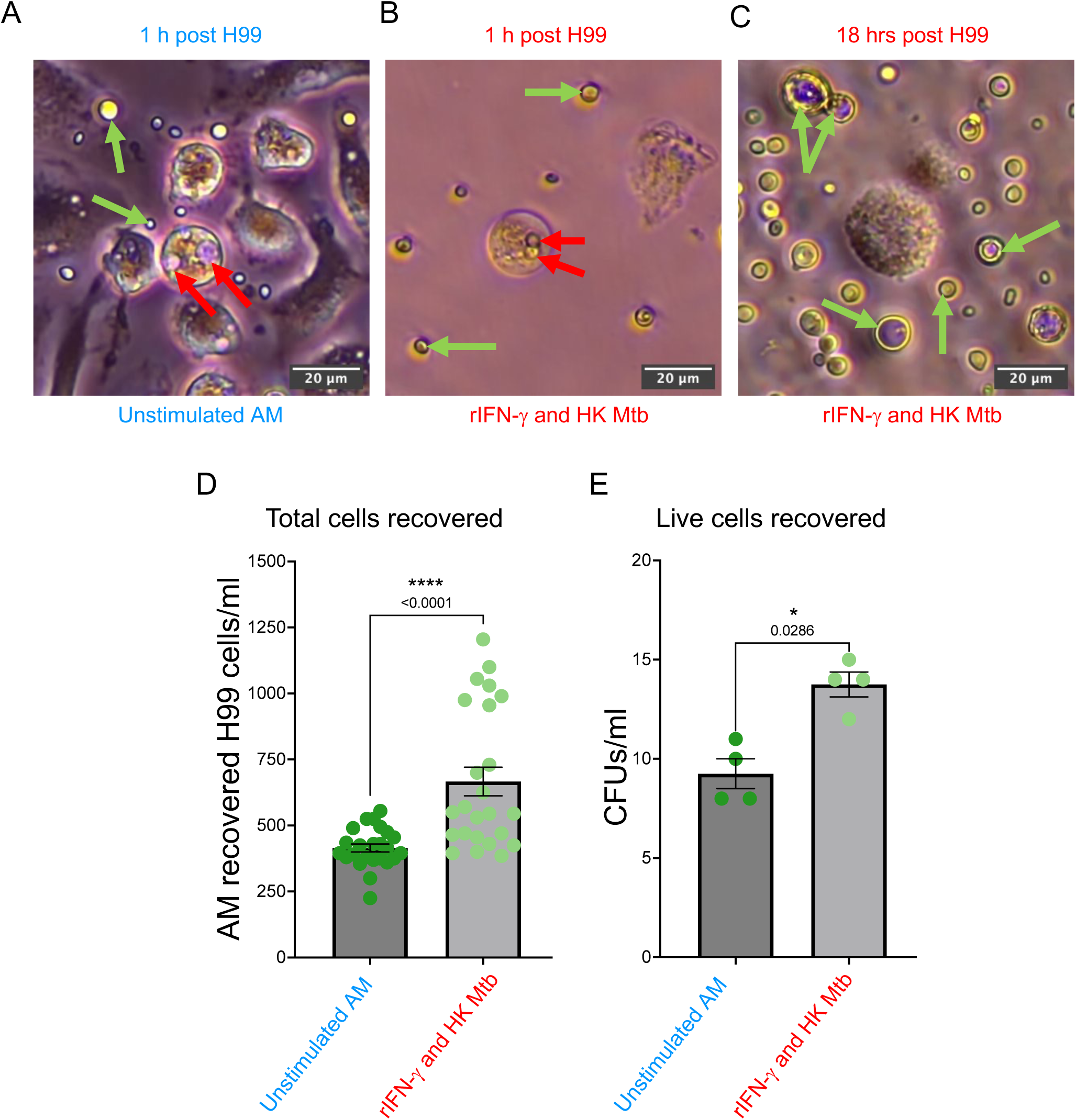
Pre-activation of alveolar-like macrophages with IFN-γ and *M. tuberculosis* cues reduces their capacity to restrict *C. neoformans* growth. Bone marrow cells from C57BL/6 mice were used to generate alveolar-like macrophages (AMs). AMs were then stimulated with heat-killed *M. tuberculosis* H37Ra and recombinant IFN-γ for 4 hours at 37°C, 5% CO_2_. 5 x 105 unstimulated and stimulated AMs/well were challenged with 2 x 10^5^ C. neoformans H99 cells at a multiplicity of infection of 5:2. AMs were imaged on an Olympus CKX53 microscope with EP50 camera, at 20x magnification with 100×100µm panels used to show; (**A**) is unstimulated AM response to 1h H99 challenge, (**B)** rIFN-γ and HK Mtb-stimulated AM response to H99, and (**C**) is stimulated AM response 18h post-H99 challenge. Red arrows indicate internalised H99 cells, green arrows indicate external H99 cells in suspension. At 18h post-infection, supernatant was removed and AMs were washed with HBSS to remove any H99 cells that were not internalised by AMs. Then, AMs were lysed with dH_2_O, and total H99 cells were quantified using a Vi-cell Blu viability analyser (**D**) or plated onto YPD agar to count live H99 cells that were internalised by AMs (**E**). Data presented are mean ± SEM from 1 biological replicate with 24 technical replicates for total cells and 4 technical replicates for CFUs. Mann-Whitney U test was used to compare *C. neoformans* cells number from unstimulated versus rIFN and HK Mtb-stimulated AMs.

## Discussion

Cryptococcosis and tuberculosis co-infections are increasingly recognized in clinical settings, particularly among individuals with compromised immunity, yet remain poorly understood and frequently underdiagnosed due to overlapping pulmonary and neurological presentations^46–49^. While the majority of available evidence derives from retrospective and post-mortem case reports, our study provides direct experimental evidence that *C. neoformans* can sense and respond to mycobacterial signals, undergoing morphological and functional changes consistent with enhanced virulence potential.

By culturing *C. neoformans* under physiologically relevant conditions^29^, we observed that exposure to either live BCG or heat-killed *M. tuberculosis* results in population-level remodelling of fungal morphology, including increased cell body size and, in the case of virulent mycobacterial stimuli, robust induction of titan cell formation. titan cells are a well-established virulence phenotype characterized by increased stress resistance, impaired phagocytic clearance, and altered immune activation^27,31,32^. That this morphotype was induced even in the absence of live mycobacterial replication suggests that *C. neoformans* is responding to conserved mycobacterial-associated molecular patterns, rather than host-derived cues alone. Notably, bacterial cell wall motifs have been shown to trigger the formation of titan cells previously^27^, supporting a model in which inter-kingdom microbial cues dynamically modulate fungal virulence traits.

Capsule remodelling was similarly enhanced across clinical isolates following mycobacterial exposure, although the direction and magnitude varied by strain. This strain-dependent variation aligns with clinical observations that differences in capsule architecture underlie distinct dissemination trajectories and immune evasion strategies^30,32,50^. Importantly, increased capsule thickness is predicted to reduce opsonophagocytic clearance^51,52^, reinforce anti-inflammatory polarization^52^, and, if such changes occur in the CNS, potentially exacerbate intracranial pressure^50^, all of which may worsen disease severity during co-infection.

The functional consequence of these morphological changes was further supported by macrophage interaction assays. When alveolar-like macrophages were primed with IFN-γ and *M. tuberculosis* components - to mimic a lung environment shaped by mycobacterial infection^41,42^ - they internalized greater numbers of *C. neoformans* without enhanced killing. Given that *C. neoformans* can survive and replicate intracellularly, particularly within alternatively activated macrophages^11,51^, this suggests that a tuberculosis-influenced immune environment may inadvertently facilitate fungal persistence. In this context, *C. neoformans* does not merely tolerate macrophage uptake but may use the intracellular niche as a protected proliferative compartment, a phenomenon consistent with previous work showing intracellular expansion and non-lytic exocytosis^11,12,51^. Together, these findings support a model in which sensing of mycobacteria by C. neoformans enhances its pathogenic potential, potentially reshaping the pulmonary immune landscape in ways that facilitate cryptococcal adaptation, persistence, and dissemination. This phenomenon parallels other examples of inter-pathogen modulation of virulence, including *Staphylococcus aureus-Candida albicans*^28,53^ and *Acinetobacter baumannii-Cryptococcus* interactions^54^, where microbial sensing promotes capsule or biofilm remodelling to enhance persistence.

## Conclusion

While this study establishes that *C. neoformans* can sense and respond to mycobacterial cues in ways that enhance virulence-associated phenotypes, several key questions remain. The specific mycobacterial components or metabolic signals responsible for initiating these morphological transitions are not yet known; identifying the molecular pathways through which *C. neoformans* detects and interprets mycobacterial presence will be essential to determine whether this response is mediated by receptor signalling, secreted factors, or direct cell-cell contact. Additionally, although our macrophage assays indicate that mycobacterial exposure primes *C. neoformans* for increased intracellular persistence, *in vivo* studies are required to determine whether these adaptations translate to altered dissemination dynamics or severity of disease during co-infection. Sequential and simultaneous infection models, particularly in lung-directed systems, will be valuable for defining how the order and timing of pathogen encounter shapes disease outcome. Finally, given that both tuberculosis and cryptococcosis co-occur in regions with limited access to advanced diagnostics, understanding whether co-infection contributes to treatment failure, delayed clearance, or relapse may have direct clinical relevance. Together, these lines of investigation will help clarify how inter-pathogen interactions shape infection trajectories and may ultimately inform therapeutic or diagnostic strategies for managing co-infection in high-burden settings.

## Materials and Methods

### Strains and Culture Conditions

*C. neoformans* and *Mycobacterium* spp., strains used in this study are summarised in **Supplementary Table 1.** *C. neoformans* clinical isolates were kindly provided by Prof. Kirsten Nielsen (Virginia Tech University, USA). Yeast cells were routinely grown on yeast extract peptone adenine dextrose (YPD) agar plates, following Chun and Madhani’s recipe^55^, stored at 4°C. For routine culture, cells were incubated at 30°C for 12-16 hours overnight in 50 mL falcon tubes with 5 mL YPD broth, recipe also by Chun and Madhani^55^, 30°C, 150RPM. Vi-cell Blu viability analyser was used to adjust for 1×10^6^ cells in 5 mL phosphate buffered saline (PBS) solution (10x, pH 7.4, Gibco™, Fisher Scientific, Loughborough, UK) for the subsequent investigations. *M. bovis* (BCG) was kindly provided for this study by Romey Shoesmith (MRC CMM, Exeter, UK). Glycerol stocks were inoculated onto Middlebrook 7H10 agar (M0303, Sigma-Aldrich, Dorset, UK), supplemented with 10% Tween-80 (655207, Sigma-Aldrich, Dorset, UK), at 37°C for 21 days. These plates were then stored at 4°C. Liquid culture was also prepared in Middlebrook 7H9 media (M0178, Sigma-Aldrich, Dorset, UK), supplemented with 10% Tween-80; 250 mL of media was added to a sterile glass conical flask aseptically. This flask was inoculated with one sterile loop of glycerol-suspended *M. bovis* stock, incubated at 30°C and 70RPM, with the media replenished weekly. A sterile conical flask was incubated with 250 mL of the same 7H9 media preparation and incubated alongside the liquid culture, to demonstrate sterility of the preparation. For *M. bovis*, OD_600_ was normalised for 1×10^6^ cells in 5 mL PBS for the subsequent investigations.

### Assessing *C. neoformans* morphological changes in the presence/absence of M. bovis *or* M. tuberculosis

50 µL of each *C. neoformans* strain inoculum were added to 12-well flat-bottomed plates, with 5 µL of *M. bovis* inoculum (seeding density of 1,000 cells/well) or 2.5ng heat-killed H37Ra *M. tuberculosis* (50ng/mL) added either at the initial inoculation or 24-hours after *C. neoformans*, with 1 mL of HPLM + 10% FBS growth media, in duplicate. Plates were incubated at 37°C in 5% CO_2_ for 48 hours, before being pelleted, washed and fixed in 4% formaldehyde in PBS for 1 hour. Cells were then spun, washed and resuspended in 50 µL PBS for imaging and further analysis.

To assess the impact on cell proliferation in the presence/absence of *M. tuberculosis*, yeast cell quantification was performed on a Vi-cell Blu viability analyser.

### India Ink staining and image acquisition

India ink staining was used to observe both *C. neoformans* cell body and capsule under the microscope. 5 µL of cell suspension was mixed with 5 µL of India Ink (Remel BactiDrop™, KS, USA) counterstain and observed under an Olympus CKX53 microscope and imaged with an Olympus EP50 camera using the Olympus EPview software (version 1.4 for Windows). These images were analysed in ImageJ2 (version 2.14.0 for macOS).

### Measurement of *C. neoformans* cell body and capsule sizes

As described above, India ink-stained cells were used to analyse *C. neoformans* cell body and capsule sizes. Cell body diameter and capsule were measured using Fiji, with two frames per sample and condition analysed, per experimental repeat (two areas per slide/four areas analysed in total). Cells were randomly selected, with *n* ≥ 200 cells total and *n* ≥ 100 cells total, per H99 condition and per clinical isolate, respectively. Capsule size was calculated by taking the cell body size from the total cell diameter measured.

### Generation of alveolar-like macrophages from murine bone marrow

C57BL/6J were purchased from Charles River Laboratories, UK. All the procedures complied with the University of Exeter ethical review process and UK Home Office licence PP9965358. Mouse bone marrow cells were isolated from wild-type C57BL/6 mice, with erythrocytes removed by RBC lysis buffer (11814389001, Sigma-Aldrich, Dorset, UK). The remaining cells filtered through a 70µm cell strainer. 5×10^5^ bone marrow cells were seeded into each well of a 12-well plate, and supplemented with DMEM (4.5g/L glucose, 12077549, Gibco™, Fisher Scientific, Loughborough, UK) containing 10% FBS, 100 U/mL penicillin-streptomycin (11548876, Gibco™, Fisher Scientific, Loughborough, UK), 20 ng/mL GM-CSF (415-ML-020, Bio-techne, Abingdon, UK) and 2 ng/mL TGF-β (7666-MB-005, Bio-techne, Abingdon, UK) (hereby referred to as GT media). These cells were maintained at 37°C, 5% CO_2_ for 7 days, with no media changes. On day 7, the media was refreshed, with the addition of 0.1 µM PPAR-γ agonist rosiglitazone (5325/10, Bio-techne, Abingdon, UK) (hereby referred to as GTR media) and incubated for a further 4 days. On day 11, nonadherent cells were washed off and discarded with Hank’s balanced salt solution (HBSS) (15266355, Gibco™, Fisher Scientific, Loughborough, UK), and re-supplemented with GTR media. On day 12, cells were detached by Accutase (A1110501, Gibco™, Fisher Scientific, Loughborough, UK) and cell-scrapers and harvested as alveolar macrophage-like cells. In total, 7.5×10^7^ AM-like cells were harvested, as quantified by Vi-cell Blu viability analyser.

### Confirmation of AM-like phenotype by flow cytometry

AM-like cells were harvested as described above, fixed with 1% formaldehyde for 30 minutes and then incubated at 4°C for 10 minutes with eFluor780 viability dye (eBioscience™, Fisher Scientific, Loughborough, UK). Then, cells were blocked by Fc-blocking buffer (anti-CD16/32, BD Biosciences, Wokingham, UK) for 10 minutes at 4°C. Cells were subsequently stained with anti-CD11b (BUV395, BD Biosciences), anti-CD11c (BV711, BD Biosciences), anti-Siglec-F (CD170 Super Bright 436, Fisher Scientific), anti-CD45 (PerCP-Cy 5.5, BD Biosciences) and anti-F4/80 (565410, BD Biosciences) at 4°C for one hour. Cells were then washed with FACS buffer containing 0.5% bovine serum albumin, 5mM EDTA with 7-AAD, before being measured by Cytek Aurora Flow Cytometer (Cytek Biosciences, Amsterdam, The Netherlands) and analysed on FlowJo (version 10.9.0 for macOS).

### Assessing the immune response to *C. neoformans* in the presence and absence of *M. tuberculosis* by alveolar-like macrophage cells

Harvested day-11 AM-like cells were seeded into four 12-well polystyrene plates at a density of 5×10^5^ cells/well and maintained in 1 mL GTR media. 2 plates were primed with 50ng H37Ra heat-killed *M. tuberculosis* (50µg/mL) and 1ng rIFN-γ (485-MI, Bio-techne, Abingdon, UK) for four hours, at 37°C, 5% CO_2_. At hour 4, all plates were inoculated with 100 µL 2×10^6^ cells/mL H99 cells, grown in HPLM, for 30 days prior to infection. These cells were filtered through 40µm cell strainers to remove any large *C. neoformans* cells that AM-like cells would be unable to internalise, and plates were incubated for 12 hours at 37°C, 5% CO_2_. To quantify internalisation of *C. neoformans*, plates were washed with HBSS to remove any suspended *C. neoformans* cells. Alveolar-like macrophages were lysed with purified water, and the resulting suspension either plated onto YPD agar plates in serial dilutions and incubated at 37°C for 48h, or yeast cells quantified using Vi-cell Blu viability analyser.

### Statistical analyses

These data were statistically analysed in GraphPad Prism 10 (version 10.0.0 for macOS). For comparisons between conditions, ANOVA analyses were carried out, alongside t-tests and Tukey’s multiple comparison tests. Statistical significance was calculated with the following classifications: * *P* ≤ 0.05, ** *P* ≤ 0.01, *** *P* ≤ 0.001, **** *P* ≤ 0.0001.

---

**Supplementary Figure 1.** The table shows the list of strains used within this study, consisting of *C. neoformans* reference strain H99, alongside four clinical isolates obtained from patients in Mulago Hospital, Kampala, Uganda or GF Jooste Hospital, Cape Town, South Africa^31^, one M. bovis reference strain (Institut Pasteur, Paris, France), and one heat-killed reference isolate of M. tuberculosis (ATCC 25177, Rockville, MD, USA). In the text, the strains are referred to indicate the location where they originate, e.g UgCl = Uganda clinical isolate or SACl = South Africa clinical isolate.

**Supplementary Table 1.**
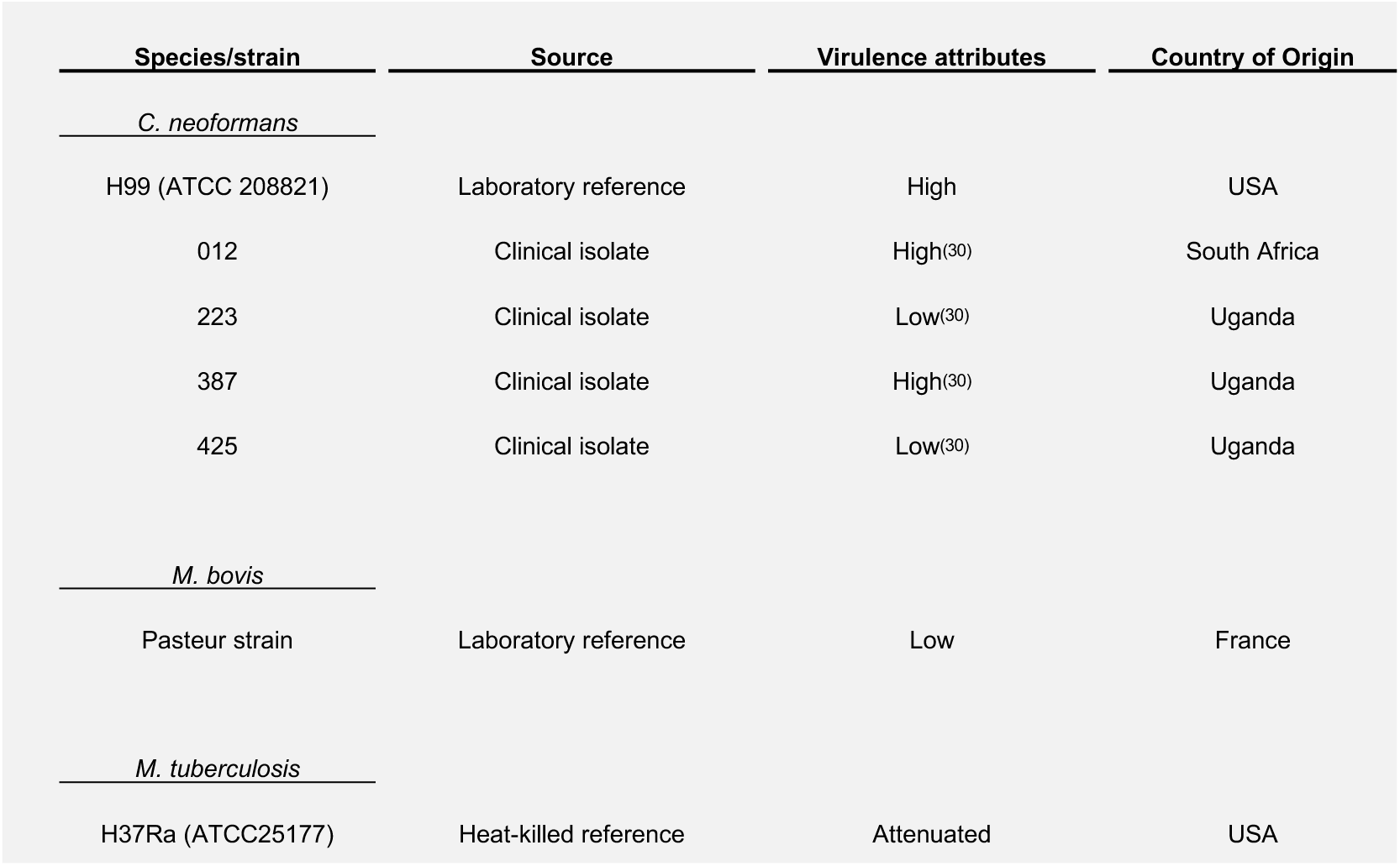
List of strains used within this study, consisting of *C. neoformans* reference strain H99, alongside four clinical isolates obtained from patients in Mulago Hospital, Kampala, Uganda or GF Jooste Hospital, Cape Town, South Africa^31^, one *M. bovis* reference strain (Institut Pasteur, Paris, France), and one heat-killed reference isolate of *M. tuberculosis* (ATCC 25177, Rockville, MD, USA).

**Supplementary Figure 2.**
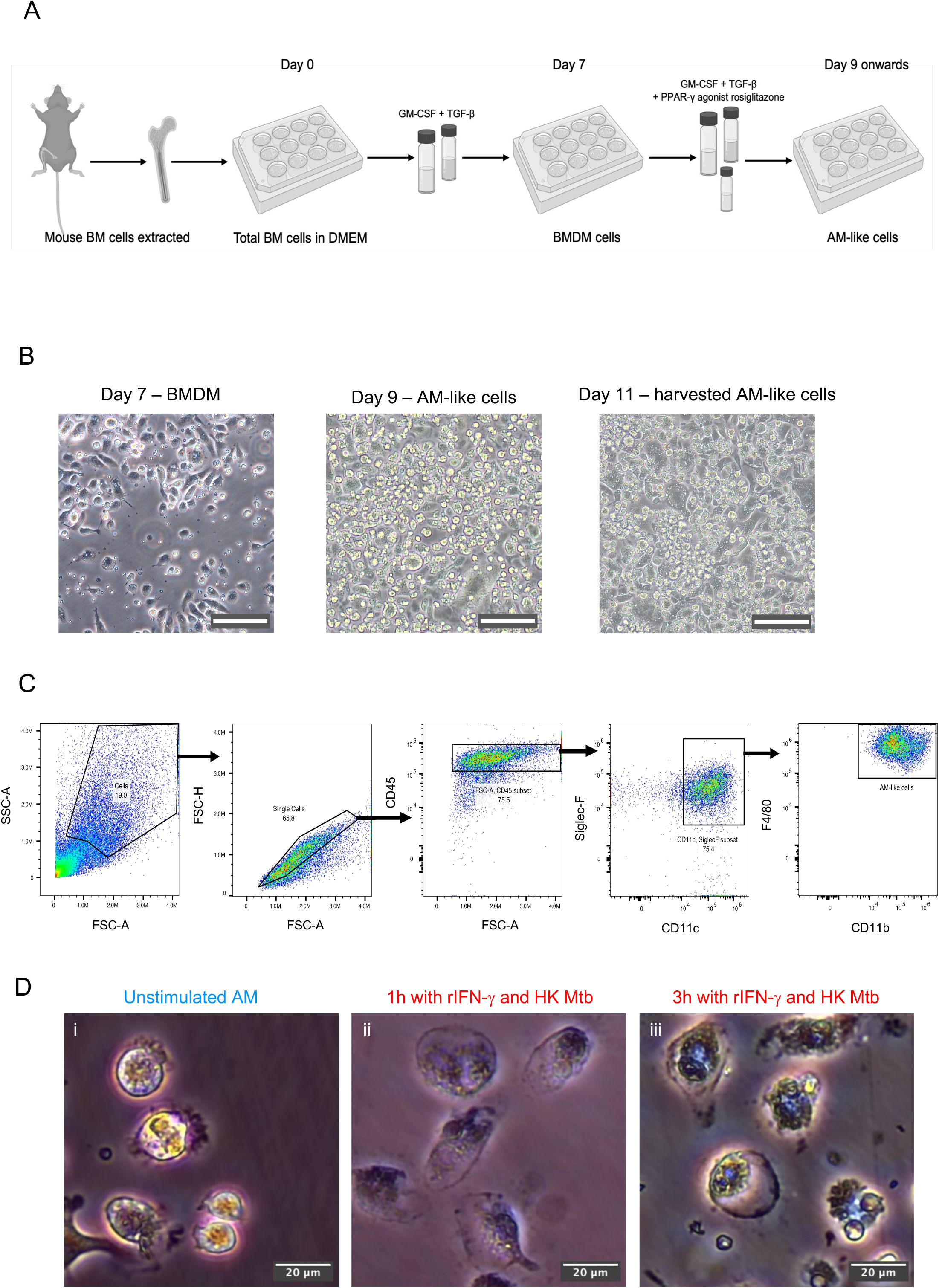
The generation of alveolar-like macrophages, validation and stimulation. Bone marrow cells from C57BL/6 mice were used to generate alveolar-like macrophages (AMs). (**A**) shows the workflow (image generated using BioRender.com). (**B**) Shows morphological changes over time during AM cell differentiation, observed on Olympus CKX53 microscope, magnification = 40x, scale bar = 50 µm. (**C**) AM differentiation was confirmed by flow cytometry staining and gating for CD11c+, Siglec-F+, F4/80high, CD11bhigh populations. AMs were then stimulated with heat-killed M. tuberculosis H37Ra and recombinant IFN-γ for 4 hours at 37°C, 5% CO_2_. (**D**) Shows AMs morphology imaged on an Olympus CKX53 microscope with EP50 camera, at 20x magnification. The 3 images represent unstimulated AMs (left), rIFN-γ and HK Mtb-stimulated AMs after 1h stimulation (centre), and rIFN-γ and HK Mtb-stimulated AMs after 3h stimulation (right).

## Acknowledgements

We thank the staff of the animal facilities at the University of Exeter for the care and support of our animals. We acknowledge funding from the MRC Centre for Medical Mycology at the University of Exeter (MR/W502649/1, MR/N006364/2 and MR/V033417/1), the NIHR Exeter Biomedical Research Centre (NIHR203320), and the Wellcome Trust (217163/Z/19/Z). The views expressed are those of the author(s) and not necessarily those of the NIHR or the Department of Health and Social Care. For the purpose of open access, the author has applied a CC BY public copyright licence to any Author Accepted Manuscript version arising from this submission. The authors declare that they have no competing interests.

